# Evaluating the Sensitivity of Sporadic Pancreatic Cancer to poly(ADP-ribose) polymerase (PARP) Inhibition (Velaparib, Olaparib, AG14361) as Single Agents and as Chemo-sensitizers

**DOI:** 10.1101/2020.03.28.013763

**Authors:** J. Angel de Soto

**Affiliations:** School of Science, Technology, Engineering and Math, Dine College, 1 Circle Dr. Tsaile, AZ 86556

**Keywords:** Pancreatic cancer, PARP1 inhibition, Olaparib, Veliparib, AG14361

## Abstract

After resection of pancreatic cancer local recurrence occurs in 50%-80% of the cases while metastasis develops 75% of the time. Current, adjuvant therapy often consists of gemcitabine, cisplatin and/or 5-fluorouracil which add a modest increase in median survival by 4-5 months. In this study, we treated human pancreatic cancer cells with poly (ADP-ribose) polymerase (PARP) Inhibitors (AG14361, Veliparib and Olaparib) alone or with gemcitabine, cisplatin or 5 – fluorouracil. Methods: CFPAC-1 and BXPC-3, HPAC human pancreatic cancer cell lines were treated for 72 hours with PARP inhibitors alone or in combination with gemcitabine, cisplatin, or 5 – fluorouracil. Validated MTT assays were used to form dose response curves from which the IC50 values were calculated. Results: The PARP1 IC50 values for CFPAC-1, BXPC-3 and HPAC pancreatic cancer cell lines were AG14361 (14.3 μM, 12.7 μM, 38.3 μM), Veliparib (52.6 μM, 100.9 μM 102.0 μM) and Olaparib (79.5 μM, 184.8 μM, 200.2 μM). The IC50 values of cisplatin, were decreased up to 60 fold in the presence of clinically relevant amounts of PARP inhibitor while 5-flourouracil IC50 values were decreased up to 6000 fold in the presence of clinically relevant amounts of PARP inhibitor. Gemcitabine was inhibited up to 73% by PARP inhibitors. Conclusions: Sporadic human pancreatic cancer cells are sensitive to PARP inhibition. PARP inhibitors significantly enhanced the cytotoxicity of cisplatin and 5-fluorouracil while inhibiting gemcitabine. There is little correlation between endogenous PARP activity and the effectiveness of PARP inhibitors.

## Introduction

Recently, Olaparib has been shown to increase progression free survival in BRCA-mutated metastatic pancreatic cancer.^1^ Adenocarcinoma of the pancreas remains a deadly disease with a dismal outcome in the overwhelming majority of cases with a 5 -year survival of only 5%.^2^ The primary therapeutic intervention in treating pancreatic cancer has been surgery with adjuvant and chemotherapy adding perhaps 2-5 months of additional overall survival when compared to surgery alone^3,4^. 5-Fluorouracil (5-FU) has had modest activity against pancreatic cancer in the adjuvant setting and in the treatment of metastatic and until recently was the mainstay of chemotherapy especially when used in combination with radiotherapy.^5.6^ Gemcitabine was shown to be able to improve median survival when used as an adjuvant over 5-FU by one month and 1-year survival by 16% over 5-FU and since, has been the mainstay of single agent adjuvant therapy.^7^ The addition of cisplatin to gemcitabine improves the median survival over gemcitabine alone by a month and a half.^8^ The efficacy of chemotherapy in pancreatic cancer clearly lags behind other diseases such as breast and colorectal cancer and current treatment protocols provide only transient relief.

Poly(ADP-ribose) polymerase (PARP) inhibitors have shown efficacy as single agents, and chemosensitizers in breast and ovarian cancer.^9^ In addition, preclinical evidence indicates that PARP inhibitors may serve as potent radiosensitizers.^10^ The PARP enzyme binds to single strand breaks (SSB) formed during base excision repair (BER) and forms complexes with pol β, XRCC1 and DNA ligase III during the repair process.^11^ The blockage of the PARP enzyme results in an eventual double strand break during DNA replication when a DNA SSB is encountered at the replication fork.^12^ These double stranded breaks may then be repaired efficiently by the process of homologous recombination (HR).^13^ In the absence of homologous recombination the cell may be repaired by the error prone nonhomologous end joining which may lead to the accumulation of chromatid breaks leading to the loss of viability.^14^ BRCA1 and BRCA2 hereditary cancers have been shown to be highly sensitive to PARP inhibitors due to their defect in homologous recombination.^15,16^ PARP inhibitors may have a broader use than treating only BRCA type hereditary cancers as a defect or deficiency in any protein (RAD51, RAD54, DSS1, RPA1, NBS1, ATR, ATM, CHK1, CHK2, FANCD2, FANCA, FANC, XRCC5, XRCC6, PNK, WRN, MRE11, RAD50, Artemis and Ligase IV) which are part of the HR pathway may render the cancer susceptible to PARP inhibition.^17^.^18^ It may be that many sporadic cancers are deficient in homologous recombination. Adding to the more generalizable use of PARP inhibitors to sporadic cancer is the finding that some keto-phenanthradine type PARP inhibitors may also cause G2/M arrest in cancer cells.^19^ It is known that cancer cells that are deficient in HR are sensitive to DNA damaging agent alone and hypersensitive to these agents after the cancer cells have been chemosensitized with PARP inhibitors.^20^ In this study we look at the use of AG14361, Veliparib and Olaparib alone or in combination with 5-FU, gemcitabine and cisplatin three of the most common agents in the treatment of pancreatic cancer.^21^ PARP inhibitors with platinum-based compounds have been approved for the treatment of BRCA deficient ovarian and breast cancers which are deficient in homologous recombination provided much of the impetus for this study.^22^ Here we expand the potential use of PARP inhibitors from BRCA mutated pancreatic cancers to sporadic pancreatic cancer.

## Materials & Methods

### Cell Lines & Medications

BXPC-3, CFPAC, HPAC human pancreatic cancer cell lines were obtained from American Type Tissue Culture in Manassas VA. 5-Fluorouracil, Cisplatin, and Gemcitabine were purchased from Sigma-Aldrich, St. Louis MO. The PARP-1 inhibitor AG14361 was obtained from Organix inc. in Woburn MA while ABT888 (Veliparib) and AZD2281 (Olaparib) were obtained from Selleck Chemical in Houston TX. Cells were grown in DMEM media with 10% fetal bovine serum in a 5% CO2 incubator at 37°C. Cells at 80-85% confluence were trypsinized, washed with PBS and plated for each experiment.

### Determining IC50 values

#### 1. Standard Curve

standard curves where made by plating 25k to 300k cells for each cell line for 4-12 hours -until the cell were attached to the wells. They were then exposed to a solution of 1 mg/ml thiazolyl blue tetrazolium (MTT) for 30 minutes. This was followed by decanting the MTT and adding propranolol to each well for 30 minutes. The absorption for each well was read with the Perkin-Elmer 1420 multi-label counter. Each data point was replicated at least 10 times.

#### 2. IC50 value

IC50 values were determined by plating 50 k cells into 12 well plates until they were attached. At this point the cells were incubated for 72 hours to various doses of drug. There were at least two controls containing vehicle for each experiment. The number of cells in each well at each specific drug concentration was then determined by the previously described MTT assay. Each experiment was replicated at least 9 times with the tenth time being a direct cell count by hemacytometer. The IC 50 values were then calculated by graphing the % inhibition vs log(dose) using Sigma Plot and 3^rd^ order equations with the ***required correlation coefficient r^2^ value having to be > 0.950***.

### Determination of PARP activity

The “PARP in vivo Pharmacodynamic Assay II kit” from Trevigen inc, Gaithersburg MD was used to determine the amount of PARP activity in each cell line prior to and after treatment with PARP inhibitors. Each assay was replicated 5 times. Endogenous Poly (ADP – ribose) in each cell line versus the IC50 if each PARP inhibitor was evaluated using the Pearson Product Moment Correlation analysis.

## Results

The ability of the PARP inhibitors AG14361, Veliparib and Olaparib to inhibit the growth of pancreatic cancer cells was evaluated utilizing the BxPC-3, CFPAC-1 and HPAC cell lines. BxPC-3 cell line was from an adenocarcinoma of the pancreas from a 61 year old female.^23^ The IC50 values derived from the dose response curves for the PARP inhibitor AG14361, Veliparib and Olaparib were 12.7 μM, 100.9 μM and 184.8 μM respectively. The CFPAC pancreatic cell line was derived from ductal adenocarcinoma from a 26 year old male with cystic fibrosis.^24^ The IC50 value for AG14361 with the CFPAC cell line was 14.3 μM very similar to that obtained for BxPC-3. CFPAC was slightly more sensitive to Veliparib and Olaparib than BxPC-3 with IC50 values of 52.6 μM and 79.5 μM. The third pancreatic cell line evaluated was the HPAC cell line another pancreatic ductal adenocarcinoma, this one was derived from a 64 year old female with liver metastasis.^25^ It was notable that these cancer cells were positive for the EGF receptor. The susceptibility to inhibition by Veliparib and Olaparib for HPAC was similar to that of the BXPC-3 cell with IC50 values of 102.0 μM and 200.2 μM respectively. HPAC however was about 3 fold less sensitive to AG14361 than either CFPAC or BxPC-3 with an IC50 value of 38.3 μM (Table 1).

**Table 1.**
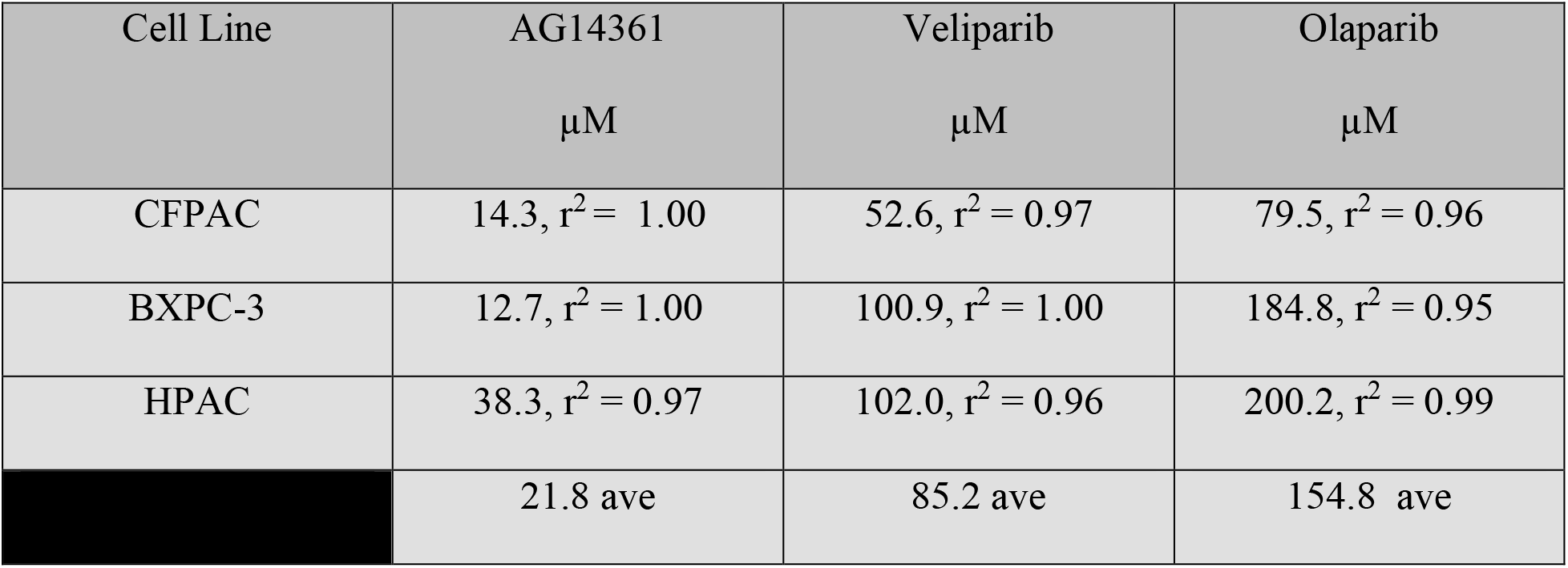
PARP inhibitor IC50 values with standard error of regression (r^2^) for sporadic pancreatic cancer

In order to better understand the significance of these results we compared the IC50 values of the preceding results with the IC50 values of BRCA1 −/− breast cancer cells. Hereditary breast cancer has been shown to be exceptionally sensitive to PARP inhibition.^26, 27^ The BRCA−/− cells examined were SUM149, HCC1937, MDA 436, and SUM 1315. The PARP IC50 values of these cell lines were compared with those of the pancreatic cell lines under identical conditions and time frames. The results were evaluated using the t test. The mean IC50 for AG14361 among the sporadic pancreatic cells was 21.8 μM as compared to 24.8 μM for the BRCA1−/− breast cancer cells with a comparative p value of 0.735. Surprisingly, the response to Olaparib was nearly identical between the pancreatic cancer cell lines and the BRCA−/− breast cancer cell lines with the mean IC50 being 155.5 μM and 155.1 μM, the p value was 0.992. Indeed, there was a trend for a greater sensitivity to Veliparib for the pancreatic cells as compared to the BRCA1−/− cells with p = 0.132 and the means of the IC50 values being 83.0 μM and 115.4 μM respectively (Table 2). After the determination of the sensitivity of sporadic pancreatic cancer to PARP inhibition was determined the next question was whether PARP inhibitors could act as chemosensitizers in pancreatic cell lines to the three most commonly used chemotherapeutic single agents to treat pancreatic cancer following surgery namely cisplatin, gemcitabine and 5-fluorouracil.

**Table 2.**
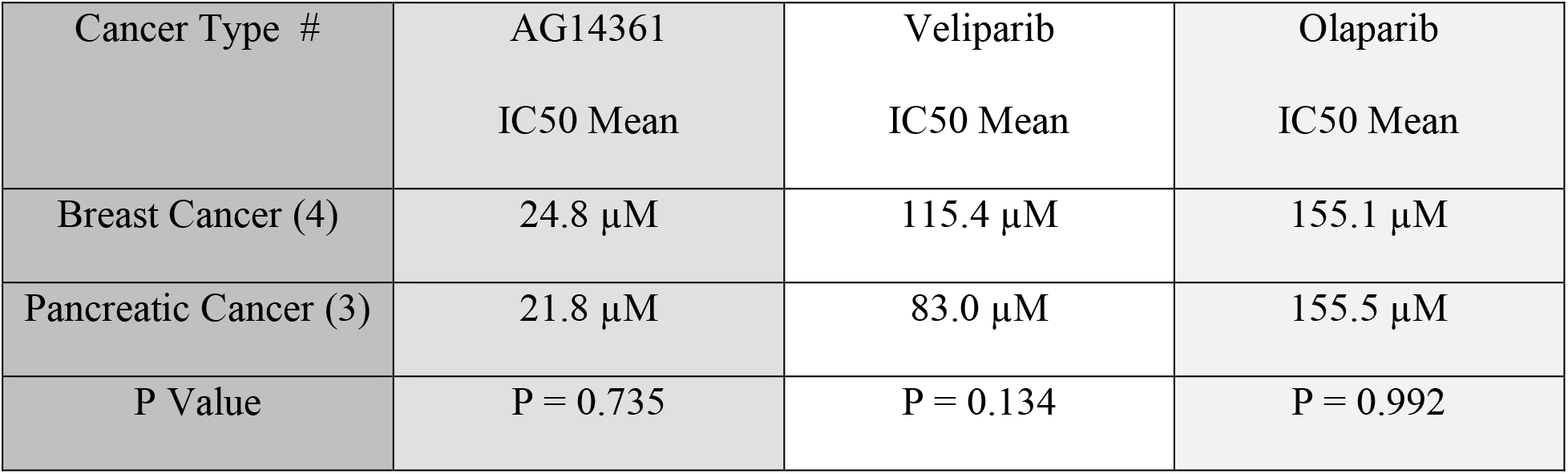
Comparative sensitivity to PARP inhibition IC50, r^2^ > 0.95

| Cancer Type # | AG14361<br>IC50 Mean | Veliparib<br>IC50 Mean | Olaparib<br>IC50 Mean |
| --- | --- | --- | --- |
| Breast Cancer (4) | 24.8 $\mu\text{M}$ | 115.4 $\mu\text{M}$ | 155.1 $\mu\text{M}$ |
| Pancreatic Cancer (3) | 21.8 $\mu\text{M}$ | 83.0 $\mu\text{M}$ | 155.5 $\mu\text{M}$ |
| P Value | P = 0.735 | P = 0.134 | P = 0.992 |

In order to keep the results clinically relevant the dose response curves for each chemotherapy agent were made in the presence of 10 μM of PARP inhibitor; that is the doses of the chemotherapy agents were varied while the concentration of PARP inhibitor was held constant at 10 μM as this may represent a reasonable upper limit of the in human *in vivo* concentrations one might reasonably achieve (calabrese; kummar;fong).^9, 28, 29^ The PARP inhibitors were able to sensitize the BXPC-3 and CFPAC cell lines to cisplatin in all cases ranging from an average increase of 63 fold with AG14361 to 1.4 fold with Olaparib (Table 3). The combination of PARP inhibitors with gemcitabine was antagonistic with the efficacy of gemcitabine being inhibited on average of 60% (Table4). 5-Fluorouracil another anti-metabolite was surprisingly enhanced by PARP inhibitors in its ability to inhibit pancreatic cancer 3100 fold by AG14361, 5.4 fold by Veliparib and 18.9 fold by Olaparib (Table 5).

**Table 3.** Chemosensitization of cisplatin by PARP inhibitors with standard error of regression (r2)

| Cell Line | Cisplatin<br>IC<sub>50</sub> | PARP Inhibitor<br>10 μM | PARP Inhibitor +<br>Cisplatin<br>IC<sub>50</sub> | Combination<br>Enhancement |
| --- | --- | --- | --- | --- |
| BXPC3 | 7.9 μM, r<sup>2</sup> = 0.99 | AG14361 | 0.13 μM, r<sup>2</sup> =<br>0.96 | **60.80 x** |
| CFPAC | 6.7 μM, r<sup>2</sup> = 0.98 | AG14361 | 0.102 μM, r<sup>2</sup> =<br>0.98 | **65.69 x** |
| BXPC3 | 7.9 μM, r<sup>2</sup> = 0.99 | Veliparib | 4.86 μM, r<sup>2</sup> =<br>0.99 | **1.63 x** |
| CFPAC | 6.7 μM, r<sup>2</sup> = 0.98 | Veliparib | 1.41 μM, r<sup>2</sup> =<br>1.00 | **4.75 x** |
| BXPC3 | 7.9, μM r<sup>2</sup> = 0.99 | Olaparib | 7.5 μM, r<sup>2</sup> = 1.00 | **1.05 x** |
| CFPAC | 6.7 μM, r<sup>2</sup> = 0.98 | Olaparib | 3.75 μM, r<sup>2</sup> =<br>0.96 | **1.79 x** |

**Table 4.** The inhibition of Gemcitabine by PARP inhibitors with standard error of regression (r2)

| Cell Line | Gemcitabine<br>IC<sub>50</sub> | PARP Inhibitor<br>10 µM | PARP Inhibitor +<br>Gemcitabine<br>IC<sub>50</sub> | Combination<br>Enhancement<br>IC<sub>50</sub> |
| --- | --- | --- | --- | --- |
| BXPC3 | 30.1 µM, r<sup>2</sup> = 0.98 | AG14361 | 112.7 µM, r<sup>2</sup> =<br>1.00 | 0.27 x |
| CFPAC | 0.46 µM, r<sup>2</sup> = 0.99 | AG14361 | 1.18 µM, r<sup>2</sup> =<br>0.95 | 0.38 x |
| BXPC3 | 30.1 µM, r<sup>2</sup> = 0.98 | Veliparib | 27.4 µM, r<sup>2</sup> =<br>0.97 | **1.09 x** |
| CFPAC | 0.46 µM, r<sup>2</sup> =<br>0.99 | Veliparib | 0.74 µM, r<sup>2</sup> =<br>0.95 | 0.62 x |
| BXPC3 | 30.1 µM, r<sup>2</sup> = 0.98 | Olaparib | 63.4 µM, r<sup>2</sup> =<br>0.95 | 0.47 x |
| CFPAC | 0.46, µM r<sup>2</sup> = 0.99 | Olaparib | 0.59 µM, r<sup>2</sup> =<br>0.96 | 0.78 x |

**Table 5.** Chemosensitization of 5-flourouracil by PARP inhibitors

| Cell Line | 5-Fluorouracil<br>IC50 ± SE | PARP Inhibitor<br>10 μM | PARP Inhibitor +<br>5-Fluorouracil<br>IC50 ± SE | Combination<br>Enhancement<br>IC50 |
| --- | --- | --- | --- | --- |
| BXPC3 | 48.6 μM, $r^2=0.98$ | AG14361 | 0.002 μM, $r^2=0.95$ | **6350 x** |
| CFPAC | 171.0 μM, $r^2=0.97$ | AG14361 | 0.50 μM, $r^2=0.95$ | **28.6 x** |
| BXPC3 | 48.6 μM, $r^2=0.98$ | Veliparib | 6.03 μM, $r^2=1.00$ | **8.05 x** |
| CFPAC | 171.0 μM, $r^2=0.97$ | Veliparib | 18.7 μM, $r^2=1.00$ | **2.81 x** |
| BXPC3 | 48.6 μM, $r^2=0.98$ | Olaparib | 1.36 μM, $r^2=1.00$ | **35.7 x** |
| CFPAC | 171.0 μM, $r^2=0.97$ | Olaparib | 81.7 μM, $r^2=1.00$ | **2.09 x** |

The endogenous activity of the poly(ADP-ribose) polymerase activity might serve as a molecular marker for determining which pancreatic cancer cells would be most sensitive to PARP inhibition. Here the IC50 values of the PARP inhibitors were correlated with the PAR levels in each pancreatic cell line. A linear regression was initially performed against PAR vs AG14361, Veliparib and Olaparib respectively with r^2^ values of 0.91, 0.9 and .08. Thus, only PAR vs AG14361 suggested any trend to a linear relationship a bit small as the slope was only a fraction being 0.32. It was observed additionally that the regression lines were all nearly horizontal indicating no correlation. A correlation analysis was performed utilizing the Pearson product moment correlation which confirmed the null hypothesis of there being no correlation (Fig 1).

**Fig 1.**
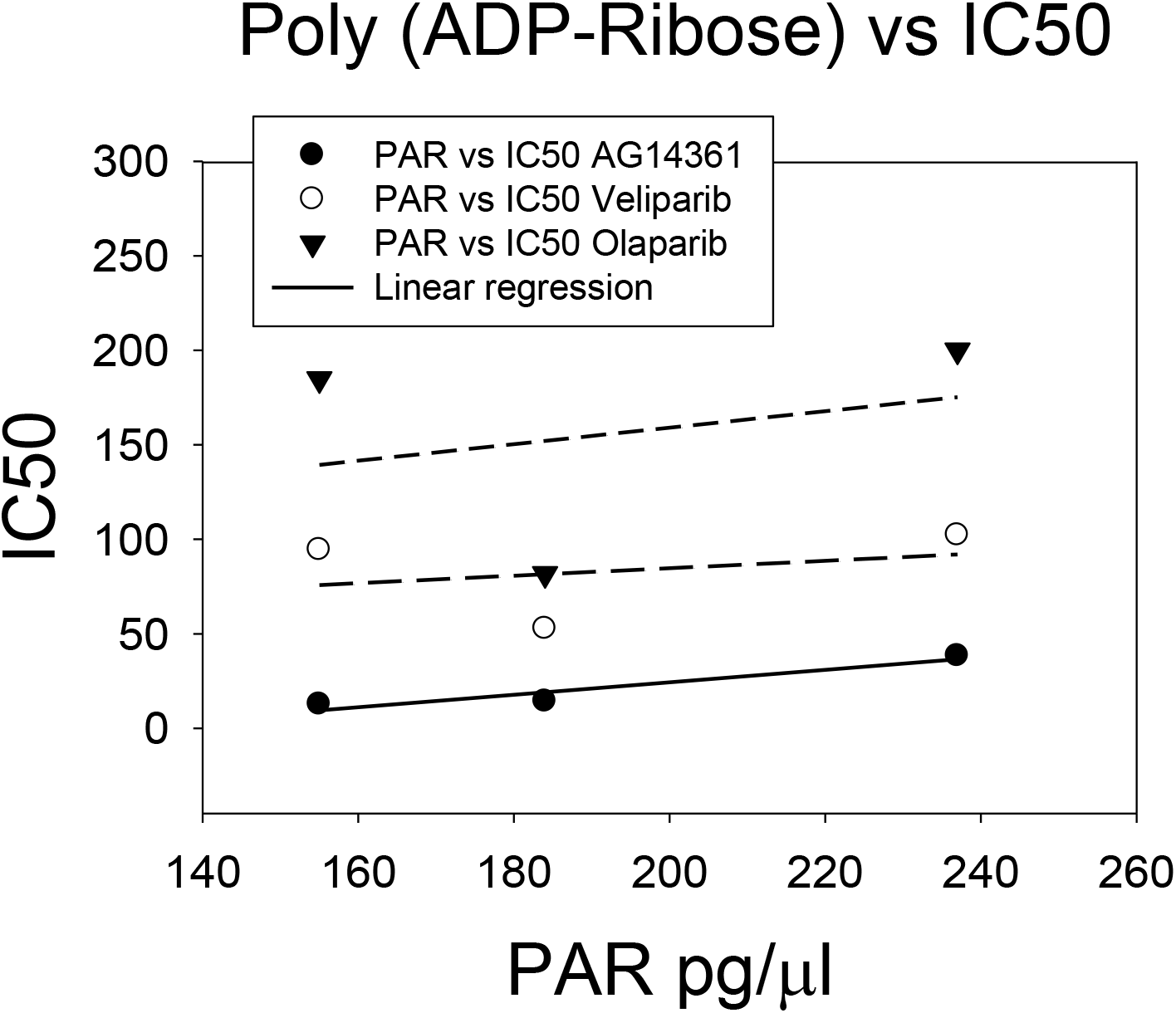
Poly(ADP-ribose) vs IC50. The IC50 values of AG14361,Veliparib and Olaparib were plotted on the y axis against the amount of endogenous poly(ADP-ribose) in each cell line prior to treatment. Both a linear regression and correlation analysis using the Pearson product moment correlation were used to evaluate the data. The p values for the Pearson coefficient were 0.191, 0.800, and 0.818 for PAR vs AG14361, Vekiparib and Olaparib respectively.

## Discussion

Clinical trials which are the gold standard in developing new therapies for pancreatic cancer often advanced without sufficient pre-clinical evidence which might be valuable in predicting the outcome of clinical trials. Some studies have for instance combined PARP inhibition with Gemcitabine even though these are clearly antagonistic in this study.^30^ The IC50 values of the three PARP inhibitors in this study were low yet the Ki for each of these three PARP inhibitors towards the PARP 1 enzyme is about 5 nM a value about ten-fold higher than used in these experiments. This implies that the inhibition of the PARP1 enzyme alone may not fully explain the inhibition of pancreatic cancer growth observed. In order to get a better idea of the potential clinical implications of the IC50 values obtained by our inhibition of sporadic pancreatic cancer cells we compared those obtained by these same PARP inhibitors against BRCA hereditary breast cancer cell line. Surprisingly, there was no statistical difference between the IC50 values of sporadic pancreatic cancer cells or hereditary BRCA1 breast cancer cells by PARP inhibitors. Indeed, inhibition by the PARP1 inhibitor Veliparib trended to be greater for sporadic pancreatic cancer as compared to BRCA hereditary breast cancer with an average IC50 for pancreatic cancer being 83 μM versus and 115.4 μM for hereditary breast cancer cells p=0.134. It is suggested that sporadic pancreatic cancers may be deficient in homologous recombination via deficiencies or defects in any of the proteins involved with this mode of DNA repair. The use of PARP inhibitors as chemo-sensitizers has long been suggested since 1996 when PARP inhibitors were used in combination with temozolomide to treat glioblastoma cell lines and later glioblastoma cancer. Here the three PARP inhibitors were used in combination with cisplatin against sporadic pancreatic cell lines. In order to make the results more translatable to a potential clinical use the PARP inhibitor concentration was held constant at 10 μM a value which could be reasonably reached in vivo while the cisplatin level was varied. The median enhancement of the inhibitory effect was 3.3 fold. Cisplatin forms both intrastrand and interstrand DNA crosslinks that induce apoptosis in the presence of mismatch repair proteins.^31^ The repair of platinum adducts is through the nucleotide excision pathway (NER) which induces SSB as obligatory intermediates thus, suggesting a role for the PARP enzyme in this repair pathway.

Experimentally, the rate of repair by the NER pathway was shown to be dependent on the PARP enzyme.^32^ We next looked at combining PARP inhibitors with the current mainstay of adjuvant pancreatic cancer treatment gemcitabine. There is evidence from breast cancer preclinical and clinical trials that PARP inhibitors enhance the efficacy of combined gemcitabine/platinum regimens.^33, 34^ Gemcitabine is metabolized to difluorodeoxycytidine di and tri phosphate where it inhibits ribonucleotide reductase and incorporates into DNA strands. In this study PARP inhibitors *antagonize* gemcitabine as a single agent in pancreatic cancer cells significantly by cutting its potency in half. The mechanism of this inhibition is unknown. 5-FU was at one time the gold standard for adjuvant chemotherapy treatment of resected pancreatic cancers. 5-FU is metabolized to fluorodeoxyuridine monophosphate which is a strong inhibitor of thymidylate synthesis additionally, in the form of the metabolite fluorodeoxyuridine triphosphate incorporates into DNA. This incorporation would induce the excision –repair process and subsequent ds DNA breaks in the presence of PARP inhibition. In the case of the BXPC-3 cell line AG14361 enhanced 5-FU ability to inhibit pancreatic cancer growth by over 6000 fold. The median enhancement of 5-FU by PARP was 22 fold. These results may suggest that 5-FU in combination with PARP inhibition has the potential to improve adjuvant chemotherapy in pancreatic cancer which up to now has been relatively disappointing. It is also significant to note that PARP inhibitors are radiosensitizers this is significant as radiation therapy is currently an important part of current treatment for pancreatic cancer.^35^

One question that has been asked is, “is there a molecular marker which one can use to predict the effect of PARP inhibition?”. The natural molecular marker of course is the PARP enzyme itself. Here we used the endogenous levels of poly(ADP-ribose) (PAR) as an indicator of PARP activity. Surprisingly there was little to no correlation between endogenous levels of PAR i.e. PARP activity and the efficacy of the PARP inhibitors against pancreatic cancer. This result was surprising as others have reported that PARP activity enhances cancer growth.^36^

## Conclusion

Sporadic pancreatic cancer cells in vitro are sensitive to PARP inhibition. PARP inhibitors significantly enhanced the cytotoxicity of cisplatin and 5-fluorouracil while <u>inhibiting gemcitabine</u>. Additional, preclinical and translational studies are needed to improve the probability of positive outcome in clinical trials.

## Acknowledgments

To the Congressionally Directed Medical Research Programs for funding this research. DOD BC grant W81XWH-07-1-0388.

